# Reactive oxygen species accelerate *de novo* acquisition of antibiotic resistance in *E. coli*

**DOI:** 10.1101/2023.07.21.550122

**Authors:** Wenxi Qi, Martijs J. Jonker, Wim de Leeuw, Stanley Brul, Benno H. ter Kuile

## Abstract

Reactive oxygen species (ROS) produced as a secondary effect of bactericidal antibiotics are hypothesized to play a role in killing bacteria. However, the role of ROS in the development of *de novo* resistance as a result of sublethal levels of bactericidal antibiotics has barely been investigated. Here, we report that single-gene knockout strains with reduced ROS scavenging exhibited enhanced ROS accumulation and more rapid acquisition of resistance when exposed to sublethal levels of bactericidal antibiotics. Consistent with this observation, the ROS scavenger thiourea in the medium decelerated resistance development. Thiourea downregulated the transcriptional level of error-prone DNA polymerase and DNA glycosylase MutM, which counters the incorporation and accumulation of 8-hydroxy-2’-deoxyguanosine (8-HOdG) in the genome. The level of 8-HOdG significantly increased following incubation with bactericidal antibiotics but decreased after treatment with the ROS scavenger thiourea. These observations suggest that in *E. coli* sublethal levels of ROS stimulate *de novo* development of resistance, providing a mechanistic basis for hormetic responses induced by antibiotics.

**Importance:** Exposure to sublethal concentrations of antimicrobials is known to result in *de novo* resistance development against the specific compound. Particularly, the use of antibiotics as feed additives to enhance productivity may result in the development of drug resistance in environmental and veterinary microorganisms, which could subsequently transfer to human populations. Nevertheless, the mechanisms underlying *de novo* resistance development have not been extensively explored. In this study, we indicate the role of ROS in promoting the formation of resistance to bactericidal antibiotics and show the potential of ROS scavengers to reduce mutation rates and slow down resistance formation under long-term selection. Thus, the supplementary use of antioxidants during prolonged antibiotic administration potentially contributes to mitigating the emergence of antimicrobial resistance.

## Introduction

Globally, the predominant proportion of antibiotic usage occurs within the realm of agricultural livestock (1)(2)(3). The utilization of antibiotics extends beyond the treatment and prevention of infections, as outside of the EU it is also employed to improve feed conversion of animals (4)(5)(6). Non-standard dosing regimens expose animals or environmental microbes to sublethal levels of antibiotics for prolonged periods of time (7)(8). These long-term sublethal levels of antimicrobials facilitate the selection of drug-resistant mutants, horizontal transfer of antimicrobial resistance genes, and *de novo* generation of drug resistance (9)(10). Resistant variants of many microbial species can easily reach humans through multiple routes, including animal-derived food products, contaminated water, soil, among others (11)(12).

Antibiotics such as quinolones, which specifically target DNA, have been observed to induce bacterial mutations partially through the activation of the SOS response (13)(14). The direct DNA damage caused by antibiotics can trigger the SOS response, leading to the upregulation of error-prone DNA polymerase genes involved in DNA repair and mutagenesis (15)(16)(17). However, it remains unclear whether exposure to other antibiotics that do not directly damage DNA, induces mutations, and facilitates development of drug resistance. Reactive oxygen species (ROS) are hypothesized to function as a secondary killing mechanism when bacteria are exposed to certain bactericidal antibiotics, like aminoglycosides, quinolones, and β-lactams (18). The specific drug-target interactions of these antibiotics generally promote the oxidation of NADH through the tricarboxylic acid (TCA) cycle-dependent electron transport chain (ETC) (19). Consequently, the formation of superoxide and hydroxyl radicals occurs, leading to damage in DNA, proteins, lipids, and the nucleotide pool, ultimately resulting in cell death (18)(20)(21)(22). Based on these observations, we hypothesized that sublethal levels of ROS may either directly or indirectly cause mutagenesis, and thereby stimulate antibiotic resistance development. Thus, depending on the concentration, ROS could be both a beneficial and a killing agent for bacteria upon exposure to antibiotics, possibly forming a mechanistic basis for described hormetic dose responses (23).

In this study, we demonstrate that *E. coli* produces ROS due to long-term exposure to sublethal levels of bactericidal antibiotics thereby facilitating *de novo* resistance acquisition. Administration of the ROS scavenger thiourea (TU) effectively reduced the mutation rate and decelerated the rate of resistance development. Additionally, the single-gene knockout strains related to ROS removal had increased levels of ROS, and their formation of drug resistance was accelerated. Notably, TU reduced lethal levels of oxidative stress, thus down-regulating the transcription of error-prone DNA polymerases induced by the SOS stress response and thereby slowing down mutation dependent resistance development.

## Results

### ROS scavenger TU reduced the rate of resistance development

To investigate the role of ROS on *de novo* acquisition of antimicrobial resistance, we exposed wild-type *E. coli* to step-increasing sub-lethal concentrations of four antibiotics and documented antibiotic resistance evolution. According to the “radical-based” theory assuming a common killing mechanism for bactericidal antimicrobials, bacteria generate ROS when exposed to bactericidal antibiotics such as amoxicillin, enrofloxacin, and kanamycin (18)(24). In contrast, the bacteriostatic antibiotics exposure generates negligible ROS in cells, so we used tetracycline as a control. Superoxide and hydroxyl radicals are scavenged efficiently by thiourea (TU). Hence, resistance development of TU-treated cultures was compared to the wild-type.

The TU-treated cultures had lower rates of resistance build-up against the bactericidal compounds than TU-absence cultures (Figure 1A-C). During amoxicillin and kanamycin exposure, differences between the wild-type and TU-added cultures were observed from the early stage, as evidenced by significantly higher minimum inhibitory concentrations (MIC) in the wild-type at day 10, day 20, and day 30 (Figure 1E and G). The most noticeable difference between the wild-type and TU-treated cultures during enrofloxacin exposure started from day 20, when the MIC of the WT-resistant strain became significantly higher than that of the TU-treated-resistant strain (Figure 1B and F). The final resistance concentrations were lower in one replicate of amoxicillin, enrofloxacin, and both replicates of kanamycin exposure in combination with TU treatment. In the case of the bacteriostatic tetracycline, the final concentration was much lower than after exposure to bactericidal antibiotics, and there were no meaningful differences between the TU-treated cells and the wild-type in resistance acquisition rate and MIC (Figure 1D and H).

**Figure 1.**
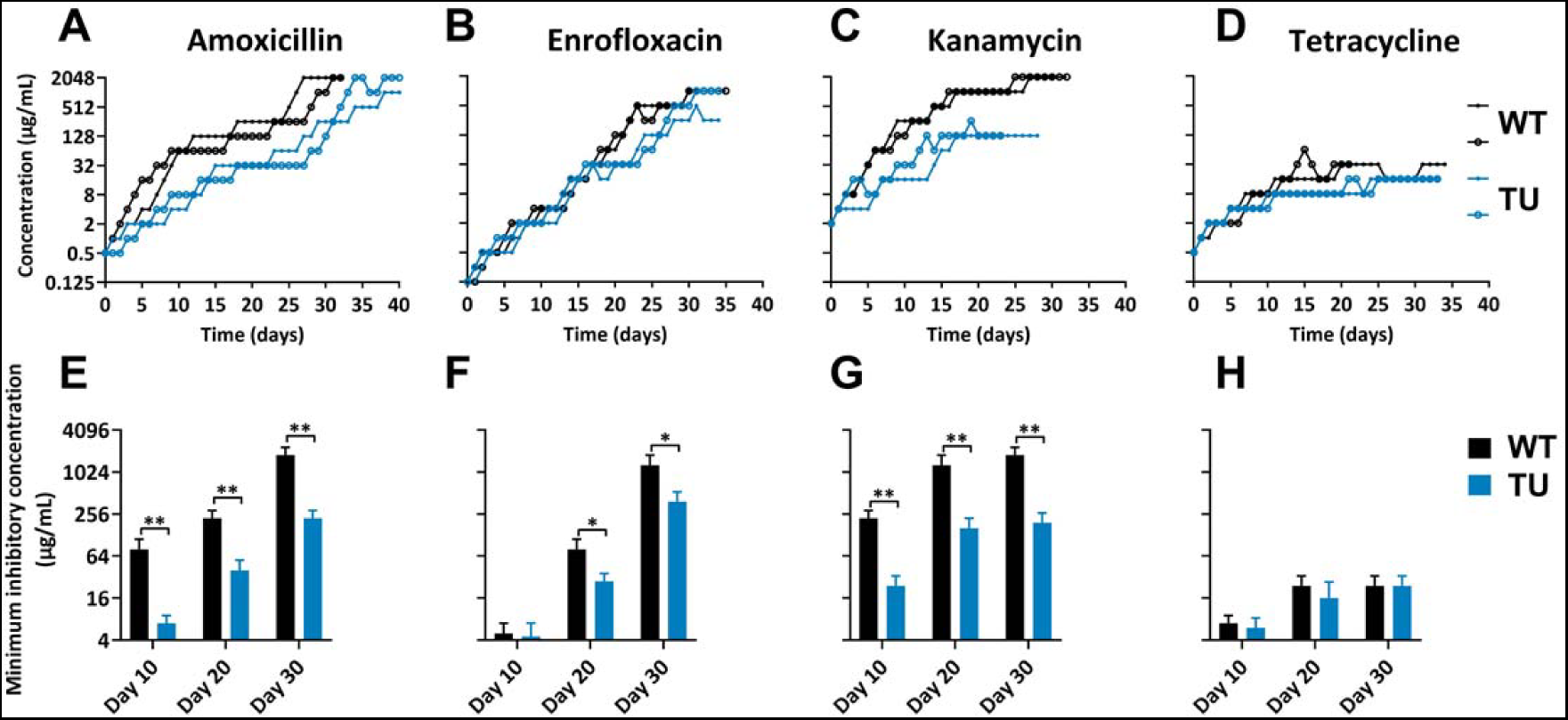
ROS scavenger TU slowed down bactericidal antibiotics resistance development. (A-D) Acquisition of resistance to amoxicillin (A), enrofloxacin (B), kanamycin (C), and tetracycline (D) of *E. coli* wild-type MG1655 (black lines) and the TU-treated (100mM) *E. coli* wild-type MG1655 (blue lines) strains. The x-axis represents the duration of evolution in days, while the y-axis represents the concentration of acquired resistance. (E-H) The minimum inhibitory concentration (MIC) at day 10, day 20, and day 30 for each resistant strain (WT and TU-treated *E. coli*) with respect to amoxicillin (E), enrofloxacin (F), kanamycin (G), and tetracycline (H) during the process of antibiotic resistance acquisition. Data are presented as means ± SD, statistical significance was determined using a one-way ANOVA, N ≥ 3, *p < 0.05, **p < 0.01.

### ROS produced during bactericidal antibiotic exposure and TU treatment

To ascertain the presence of ROS at the moment that the cell was exposed to increased levels of an antibiotic, the intracellular levels of ROS were visualized using widefield fluorescent microscopy. Cells were taken from cultures at the middle stages where the MIC increased rapidly when the cells reached the resistance concentrations of 32 μg/mL. Time-lapse images showed that the intracellular presence of ROS was low immediately at the start of the exposure to one of the bactericidal antibiotics, increased after 3-5 hours, and decreased again approximately two hours later (Figure 2A-C). ROS production levels were highest at 6 hours of amoxicillin or enrofloxacin exposure, or 4.5 hours of kanamycin exposure. Hardly any ROS was observed in the tetracycline-exposed strain (Figure 2D). Subsequently, ROS production was quantified by flow cytometry in these strains exposed to 32 μg/mL amoxicillin, enrofloxacin, kanamycin, and tetracycline (Figure 2E). The bactericidal antibiotics induced higher ROS levels than the bacteriostatic tetracycline. TU treatment significantly reduced the ROS levels caused by exposure to bactericidal antibiotics. Flow cytometry sorting was applied to divide the ROS positive and negative populations in these strains when ROS levels were highest, and the DNA of the separated populations was sequenced entirely. No different mutations between the ROS positive and negative populations were observed (Table S1).

**Figure 2.**
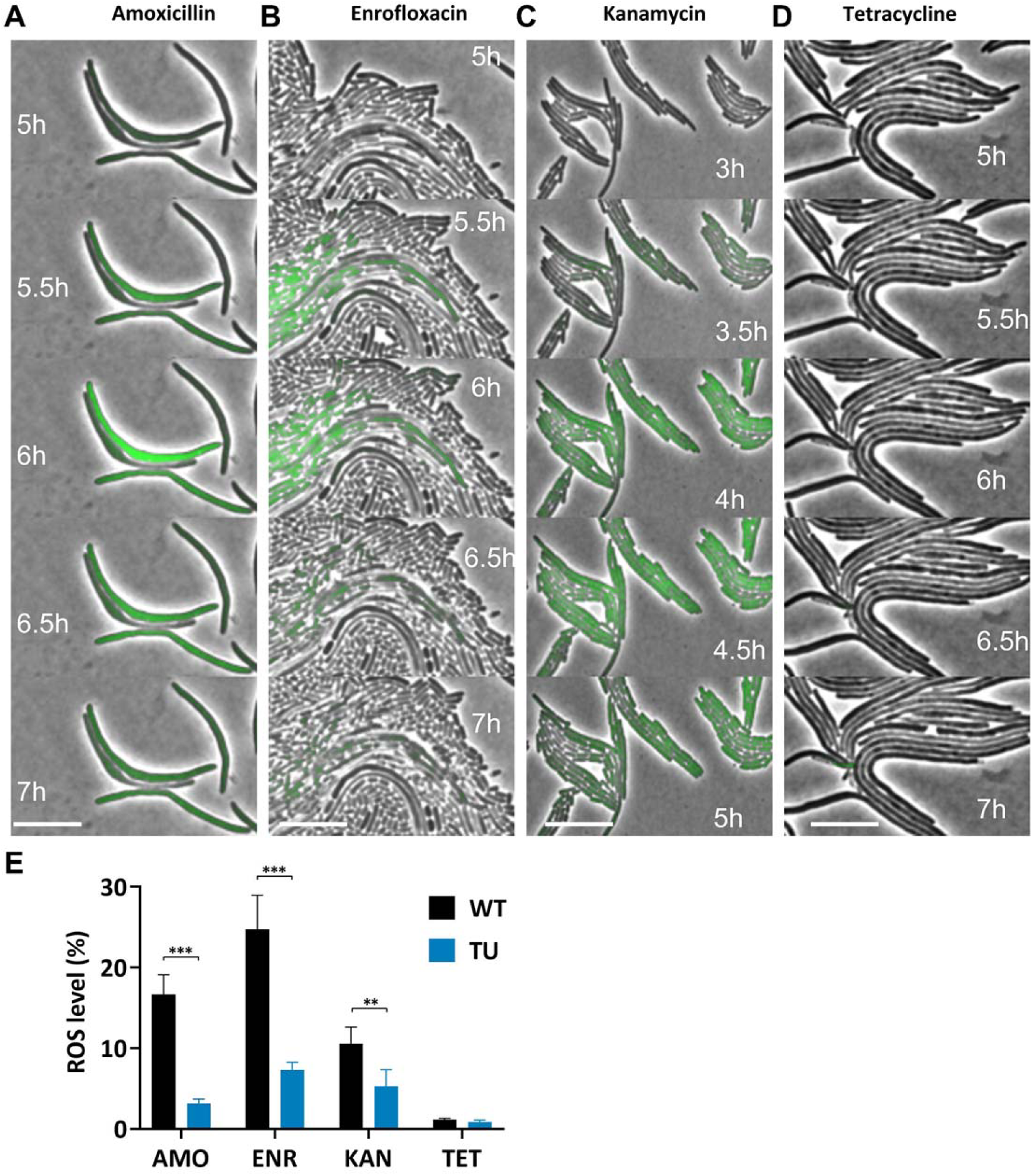
ROS produced during bactericidal antibiotic exposure and attenuated by TU treatment. (A-D) Time-lapse photography capturing the ROS generation in *E. coli* exposed to antibiotics treatment with cells resistant to 32 μg/mL amoxicillin (A), enrofloxacin (B), kanamycin (C), and tetracycline (D). The scale bar refers to 10 μm. (E) ROS production levels were quantified by flow cytometry. *E. coli* and TU-treated *E. coli* resistant strains were exposed to 32 μg/mL of amoxicillin (AMO), enrofloxacin (ENR), kanamycin (KAN), and tetracycline (TET), separately. The strains used are the same as A-D. Data are represented as means ± SD, statistical significance was investigated using a one-way ANOVA, N = 3, **p < 0.01, ***p < 0.001.

### ROS elimination-associated single-gene knockout strains acquire resistance faster

While the previous set of experiments examined the effects of reduced ROS on the development of resistance, the effect of increased ROS was investigated using six single-gene knockout *E. coli* strains, Δ*sodA*, Δ*sodB*, Δ*soxR*, Δ*soxS*, Δ*katE*, and Δ*yggX* that are deficient in ROS elimination. SodA and SodB are superoxide dismutase (25). SoxR activates the transcription of SoxS, and both SoxR and SoxS participate in the removal of superoxide (26). The KatE enzyme is the monofunctional catalase, which decomposes hydrogen peroxide into water and oxygen (27). YggX is a putative Fe^2+^-trafficking protein, which is proposed to play a role in the oxidation resistance of iron-sulfur clusters (28). We assume that when these genes are removed or disabled, bacteria will lose part of the ability to remove excess ROS, thus increasing the ROS levels during antibiotic treatment.

After prolonged amoxicillin exposure, maximum resistance concentrations of the mutants and wild-type strains were similar (Figure 3A). However, the rates of resistance development in these single-gene knockout strains were faster than the wild-type during the middle-late stages, as evidenced by significantly higher MIC in Δ*sodA*, Δ*soxR*, and Δ*soxS* at day 20 (Figure 3E). For enrofloxacin, the MIC of the knockout strains started to increase after day 10. Both Δ*soxR* and Δ*yggX* exhibited significantly higher MIC compared to the wild-type at day 10 and day 20, while the MIC of Δ*sodA* and Δ*katE* was also higher than the WT-resistant strain at day 20 (Figure 3B and F). The final enrofloxacin resistance of the knockout-mutant strains reached maximum concentrations of 2048 μg/mL, double compared to the wild-type (Figure 3B). Of all knockout-mutant strains except Δ*soxS* only one replicate could reach the maximum concentration. The other replicates were killed while evolving to resist to high concentrations of the antibiotic, probably by the elevated ROS levels. Despite this, the surviving replicates in Δ*sodA*, Δ*soxR*, Δ*katE*, and Δ*yggX* showed faster resistance acquisition corroborating that ROS at these levels lead to cellular conditions that are at a tipping point between being beneficial and detrimental. During kanamycin exposure, all mutant strains reached the same maximum resistance concentrations as the wild-type strains (Figure 3C). However, mutant strains reached 2048 μg/mL faster than the wild-type, and the MIC of Δ*sodA* and Δ*soxR* were significantly higher than the wild-type at day 10 (Figure 3G). No notable differences were observed in resistance development during tetracycline exposure, only in Δ*yggX* had a higher MIC than the wild-type, others, as the speed and MIC were roughly the same as the wild-type (Figure 3D and H).

**Figure 3.**
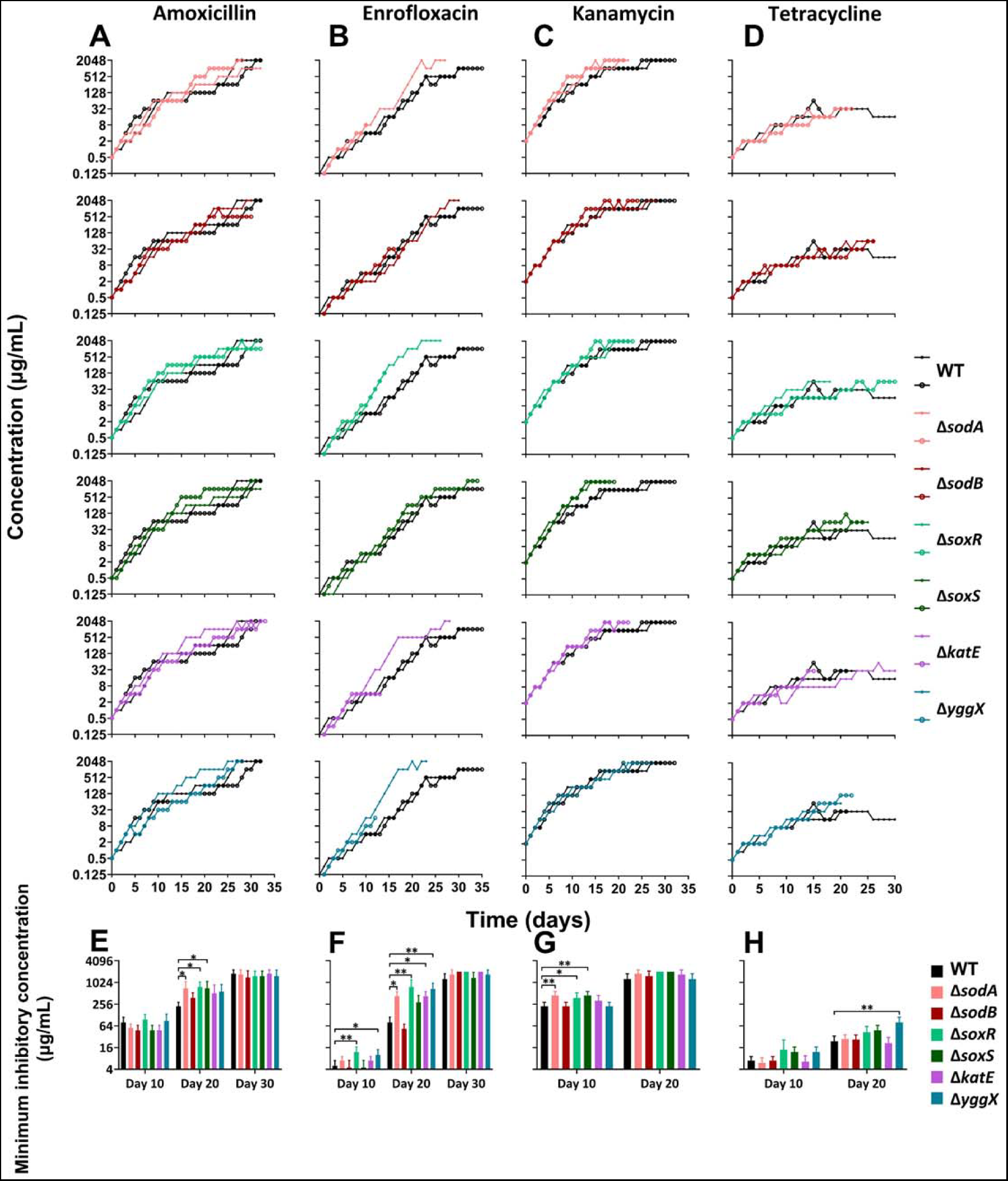
ROS elimination-associated single-gene knockout strains gain antibiotic resistance faster. (A-D) Acquisition of resistance to amoxicillin (A), enrofloxacin (B), kanamycin (C), and tetracycline (D) of *E. coli* wild-type MG1655 (black lines) and the single-gene knockout strains Δ*sodA*, Δ*sodB*, Δ*soxR*, Δ*soxS*, Δ*katE*, Δ*yggX* (colored lines). The x-axis represents the duration of evolution in days, while the y-axis represents the concentration of acquired resistance. (E-H) The MIC at day 10, day 20, and day 30 for each resistant strain (WT, Δ*sodA*, Δ*sodB*, Δ*soxR*, Δ*soxS*, Δ*katE*, and Δ*yggX E. coli*) with respect to amoxicillin (E), enrofloxacin (F), kanamycin (G), and tetracycline (H) during the process of antibiotic resistance acquisition. Data are presented as means ± SD, statistical significance was determined using a one-way ANOVA, N ≥ 3, *p < 0.05, **p < 0.01.

### ROS level elevated after knockout of *sodA* or *soxR* during exposure to bactericidal antibiotics

Subsequently, we measured the ROS production levels under two conditions. Before inducing resistance, the naïve strains were treated with one-quarter MIC antibiotics, and after acquiring *de novo* resistance, the final resistant strains were treated with the maximum resistance concentrations of these four antibiotics. We chose two single-gene knockout strains Δ*sodA* and Δ*soxR* that acquired resistance faster compared to the wild-type during exposure to bactericidal antibiotics (Figure 3). In addition, the wild-type and TU-treated wild-type were subjected to the measurement as well. The antibiotic induced ROS production levels were lower in naïve wild-type strains compared to in the *de novo* resistant strains (Figure 4A and B). TU treatment significantly decreased the ROS levels during amoxicillin and enrofloxacin exposure (Figure 4A). ROS production was significantly increased in Δ*sodA* during enrofloxacin and kanamycin exposure, and in Δ*soxR* during enrofloxacin exposure (Figure 4A). Higher but still non-lethal ROS production was detected in final resistant mutant strains (Figure 4B). TU treatment decreased ROS levels, compared to the wild-type strains, Δ*sodA*, and Δ*soxR* significantly increased the ROS production levels during exposure of the maximal bactericidal antibiotics (Figure 4B). The ROS production levels in the naïve strains or the *de novo* resistant strains were low during tetracycline treatment (Figure 4A and B). In short, when ROS-scavenger genes were knocked out, the rate of antibiotic resistance acquisition was higher, accompanied by increased ROS accumulation levels. Together with the TU-treated condition, we conclude that sublethal ROS-increase in cells correlated to enhanced antibiotic resistance development.

**Figure 4.**
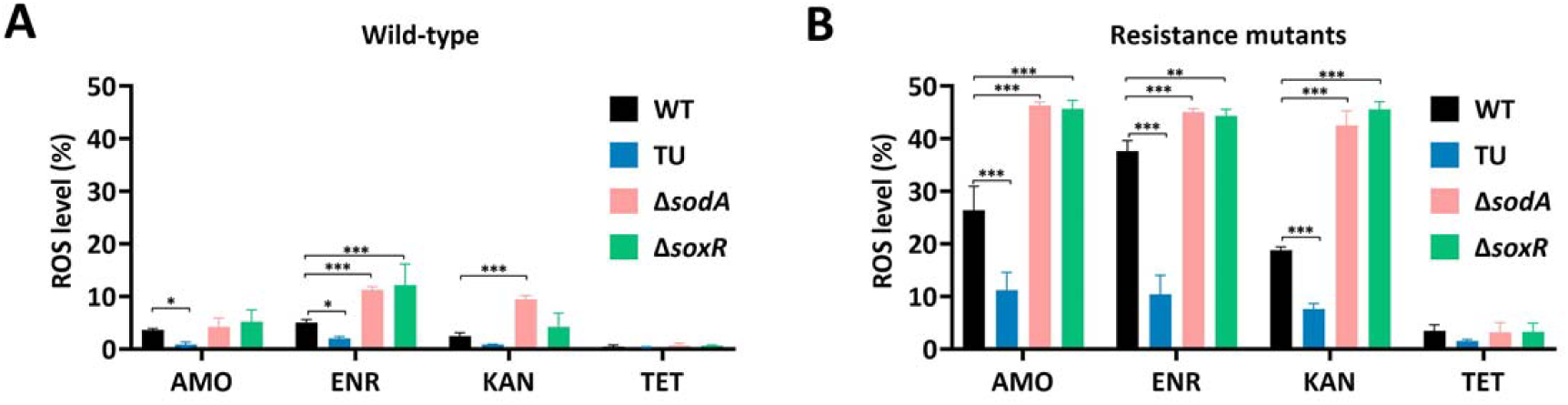
ROS level elevated after knockout of *sodA* or *soxR* during exposure to bactericidal antibiotics. (A) ROS production levels quantified by flow cytometry in naïve wild-type strain, TU-treated strain, single-gene knockout strains Δ*sodA* and Δ*soxR*, exposed to one-quarter MIC of amoxicillin, enrofloxacin, kanamycin, and tetracycline. Data are presented as means ± SD, statistical significance was determined using a one-way ANOVA, N = 3, *p < 0.05, **p < 0.01 ***p < 0.001. (B) ROS production levels quantified by flow cytometry. The final resistant strains after resistance evolution, WT-resistant strain, TU-resistant strain, single-gene knockout-resistant strains Δ*sodA* and Δ*soxR*, exposed to the maximum concentrations of amoxicillin, enrofloxacin, kanamycin, and tetracycline. Data are presented as means ± SD, statistical significance was determined using a one-way ANOVA, N = 3, *p < 0.05, **p < 0.01 ***p < 0.001.

### TU decreased the mutation rate during exposure to bactericidal antibiotics

Acquisition of antibiotic resistance is known to be accompanied by the accumulation of mutations (29). To examine whether ROS is related to mutation formation, we performed fluctuation assays to check the spontaneous mutation rate in different ROS-producing strains. The *E. coli* wild-type, TU-treated wild-type, Δ*sodA*, and Δ*soxR* naïve strains were treated with one-quarter MIC of the four antibiotics used throughout this study. 50 μg/mL rifampicin LB plates were used to select the mutated cells. The mutation rate was significantly increased after enrofloxacin and kanamycin exposure in the wild-type and in Δ*soxR* strains compared with the untreated control (Figure 5). Kanamycin exposure also significantly increased the mutation rate in Δ*sodA*. Compared with the wild-type, TU treatment significantly decreased the mutation rate during exposure to the bactericidal antibiotics amoxicillin, enrofloxacin, and kanamycin. Tetracycline exposure did not influence the mutation rate. In summary, the spontaneous mutation rate was related to ROS production, and TU treatment decreased the mutation rate.

**Figure 5.**
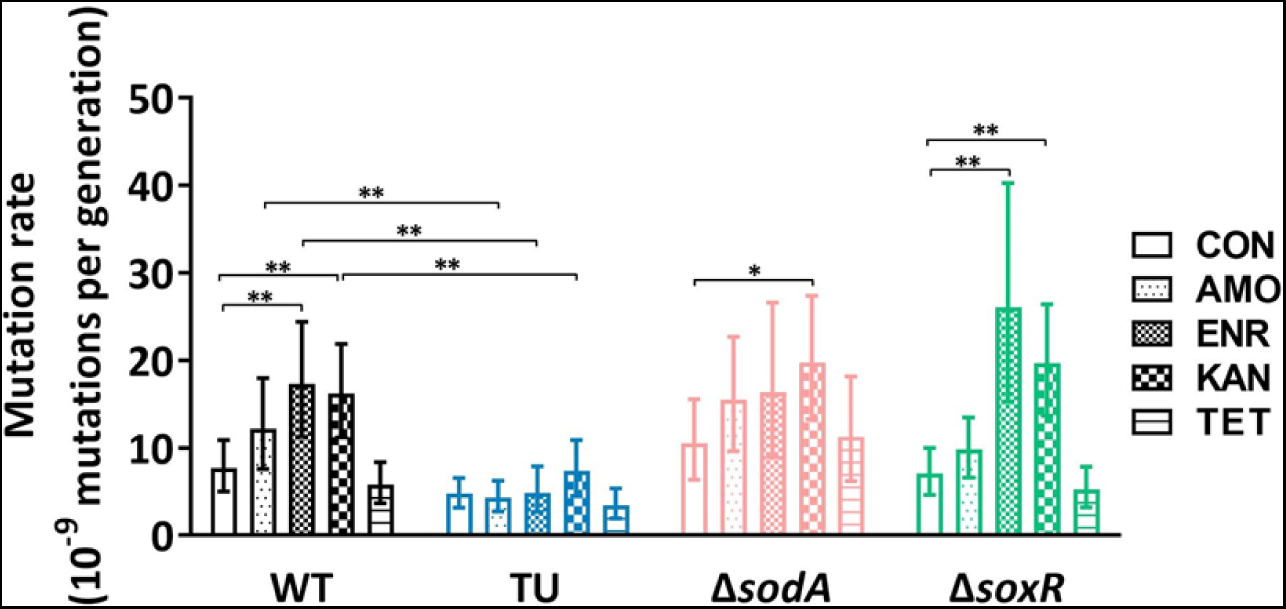
Effect of TU on the mutation rate during exposure to antibiotics. Mutation rates were measured by fluctuation analysis. The naïve strains of wild-type (WT), TU-treated wild-type (TU), Δ*sodA* and Δ*soxR*, exposed to one-quarter MIC of amoxicillin (AMO), enrofloxacin (ENR), kanamycin (KAN), and tetracycline (TET). Cells were plated on rifampicin (50 µg/mL) LB plates for counting mutated cells, and LB plates for counting the total cell number. The CFU was counted after 48 hours and analyzed by webSlvador (30). Mutation rates were calculated by Lea-Coulson Ɛ < 1.0, the comparison was done by the maximum likelihood ratio statistical test. Error bar represented the upper and lower limit of the 95% confidence intervals, *p < 0.05, **p < 0.01, N = 10, CON = untreated control culture.

### Mutated genes correlate with antibiotic resistance, oxidative stress, and SOS response

The main antimicrobial resistance mechanisms by which genes are mutated after amoxicillin exposure are antibiotic inactivation, target alteration, efflux pumps, and reduced permeability (Figure 6A). All amoxicillin-exposed strains contained mutations in *ampC* and *ompC*. AmpC is a β-lactamase with substrate specificity for amoxicillin. The mutations observed in *ampC* were located in the promoter area (Table S2). These mutations could trigger the *ampC* amplifications observed before (31). Each *de novo* resistant strain carried an *ampC* amplification contig, except one replicate in Δ*katE.* These contigs had different lengths and copy numbers but they all affected the degradation of amoxicillin (Table 1). OmpC is an outer membrane porin, mediates the entry of various antibiotics including β-lactams (32). Another frequent mutation is *envZ*, which is the sensor histidine kinase of the EnvZ/OmpR two-component system, regulates OmpF and OmpC to modulate osmosis (33). In Δ*katE* and Δ*yggX* strains, an excision of prophage element *e14* was found, which is known to occur upon SOS response activation (34).

**Figure 6.**
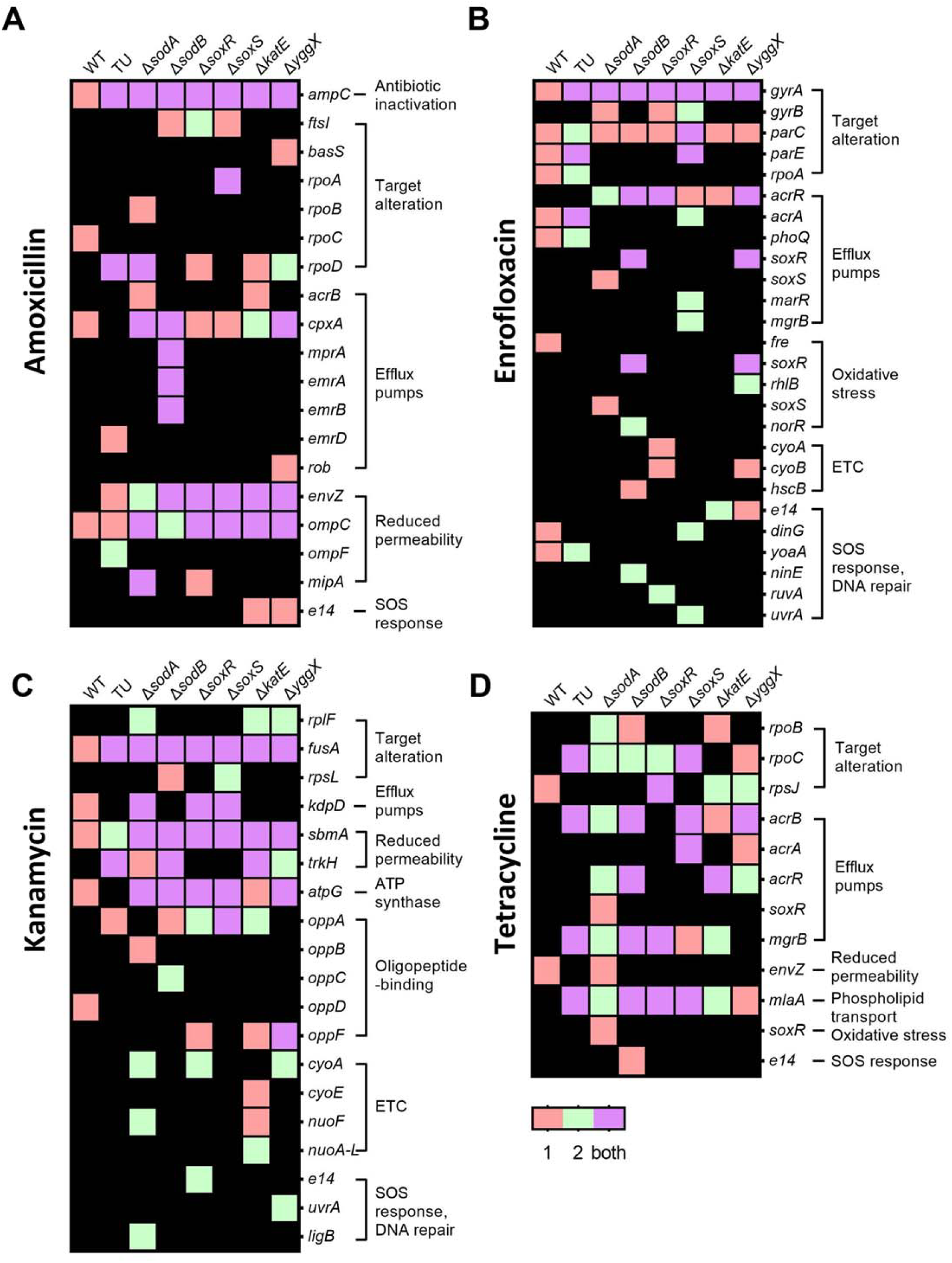
Mutated genes involved in antibiotic resistance, oxidative stress, and SOS response. (A-D) The genomic DNA sequencing of each final resistant strain compared to each no-antibiotic-treated strain to identify mutated genes. The mutated genes are determined as defined by the Comprehensive Antibiotic Resistance Database (CARD). Rows show genes that observed mutations, columns indicate different strains. Genes are grouped by functions. Red color indicates mutations observed in replicate one, green color mutations found in another replicate, purple color means mutations observed in both replicates, Black background indicates an absence of mutations. (A) amoxicillin exposure, (B) enrofloxacin exposure, (C) kanamycin exposure, (D) tetracycline exposure.

**Table 1.**
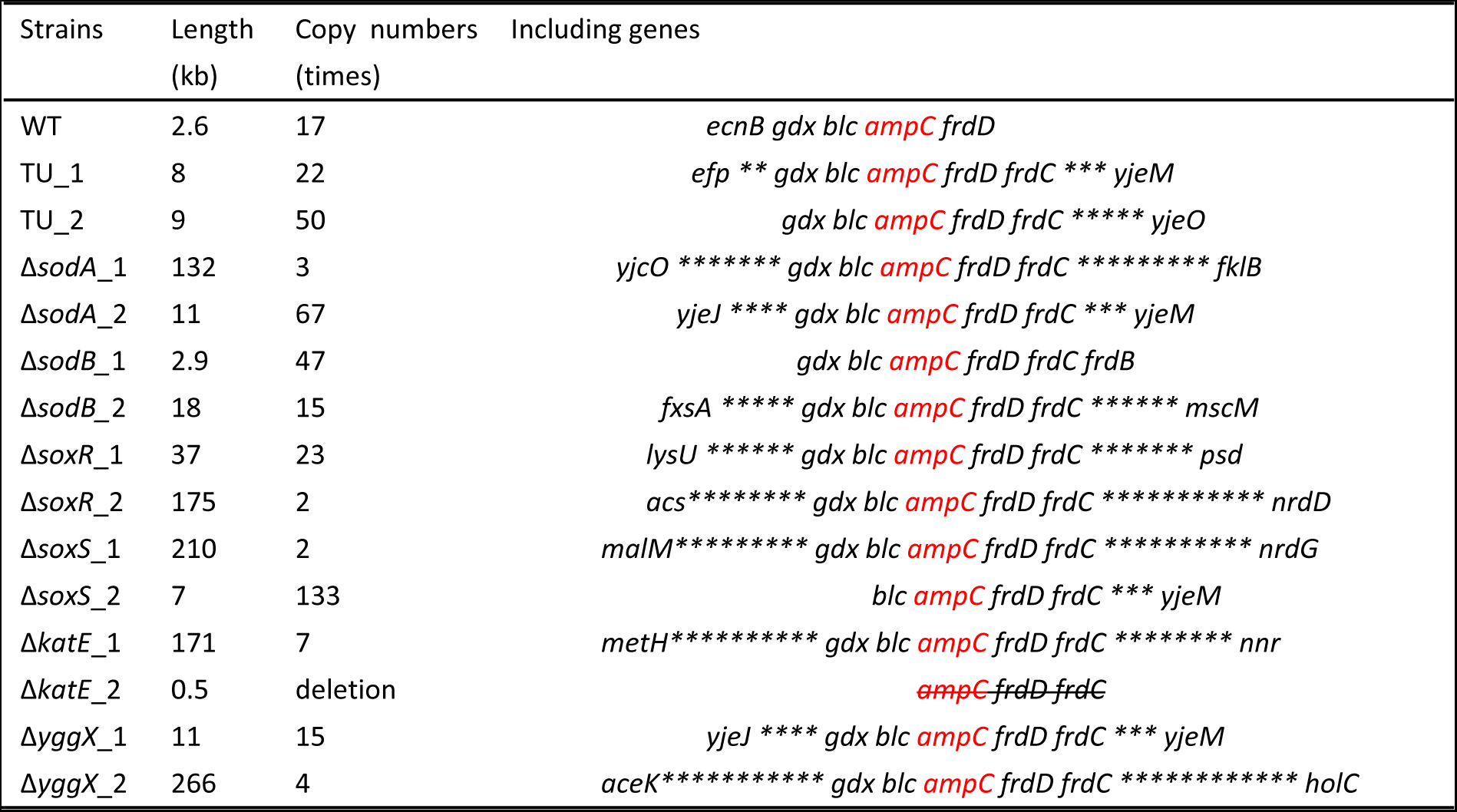
Copy numbers of contigs including *ampC* after amoxicillin exposure.

Exposure to enrofloxacin induced the mutations in *gyrA* that are usually involved in *de novo* enrofloxacin resistance (35). Similar mutations were evolved in the genes coding for gyrase subunit *gyrB*, DNA topoisomerase 4 subunit *parC*, and *parE* (Figure 6B). The single-gene knockout strains accumulated mutations in HTH-type transcriptional regulator *acrR*, which regulates *acrAB* genes. In the wild-type and the TU-treated wild-type, mutations in *acrA* were observed, which codes for a multidrug efflux pump that uses the electrochemical proton gradient to export antibiotics (36). Several mutations were observed in genes associated with oxidative stress response, the electron transport chain (ETC), SOS response, and DNA repair, especially in the single-gene knockout strains. This indicates that ROS is involved in resistance mutation acquisition.

The frequently mutated genes after kanamycin exposure are *fusA*, *kdpD*, *sbmA*, *trkH*, and *atpG* (Figure 6C). Mutations in *atpG* and *oppABCDF* have not yet been recorded in the Comprehensive Antibiotic Resistance Database (CARD), even though these mutations emerged after aminoglycosides antibiotics exposure (37) (38). Comparable to enrofloxacin-exposed strains, kanamycin-exposed strains developed mutations in genes that are involved in ETC, SOS response, and DNA repair. This further strengthens the notion that ROS is involved in the development of *de novo* drug resistance.

Tetracycline exposure caused the fewest mutations (Figure 6D). The mutations that occurred caused target alteration or appeared in genes involved in coding for efflux pumps. The genes mutated after tetracycline exposure were sometimes also mutated after exposure to other antimicrobials, albeit in different positions (Table S2). For instance, the gene coding for RNA polymerase subunit *rpoBC* was mutated also after amoxicillin treatment in wild-type *E. coli* and in the Δ*sodA* mutant (Figure 6A). The multidrug efflux pump subunit AcrA and regulator AcrR have been shared with enrofloxacin-resistant strains (Figure 6B). The *mlaA* gene that was mutated in most tetracycline-exposed strains is closely associated with *ompC* (39).

### TU modulates SOS-response genes, reducing *mutM* transcription as well as 8-HOdG levels after bactericidal antibiotic exposure

“Radical-based” theory suggests that ROS has a secondary role in cell-killing by damaging DNA and nucleotide pools. However, cells have damage-repair mechanisms such as base-excision repair and mismatch repair. Incorrect repair causes mutations in surviving bacteria, eventually promoting also beneficial mutations and thus resistance acquisition. To check transcription levels of DNA damage-repair-related genes, we performed RNA sequencing on cells exposed to antimicrobials. During amoxicillin exposure, the transcription of the DNA damage-inducible genes of the SOS response *lexA*, *umuC*, *umuD*, *mutM*, *yebG*, and *nudG* was upregulated. The increase in the transcription level was attenuated when TU was added (Figure 7A). Transcript levels of *recA*, *recN*, and *recX* were similarly upregulated, addition of TU made no difference. The up– or down-regulation of genes related to DNA damage repair showed the greatest change during enrofloxacin exposure. Compared with the TU-treated group, only *recF*, *uvrA*, *mutM*, and *vsr* displayed clearly higher expression levels in the group without TU. Most of the other genes, like *dinB*, *dinI*, *recA*, *recN*, *recX* were upregulated to similar levels with or without TU addition. After kanamycin exposure, transcription levels of *umuC*, *umuD*, and *mutM* in the wild-type were higher than those in the TU-treated group. Treatment with the bacteriostatic tetracycline showed the lowest transcription levels, and no clear difference due to the addition or absence of TU. Interestingly, the gene whose transcription level was always upregulated after bactericidal antibiotic treatment and whose transcription level was higher in the absence of TU was *mutM*. MutM is involved in base excision repair of DNA damaged by the mutagenic lesion 8-hydroxy-2’-deoxyguanosine (8-HOdG), and other oxidized nucleobase damage (40). On the one hand, the upregulation of base excision repair gene *mutM* in the TU absence group indicates that cells faced more oxidative stress during bactericidal drugs exposure, which suggests a higher mutagenesis frequency compared to the TU-treated cells. On the other hand, TU treatment decreased the mutagenesis caused by oxidative damage thus presumably slowing down mutagenesis and, also, lowering the changes of acquiring beneficial stress related mutations.

**Figure 7.**
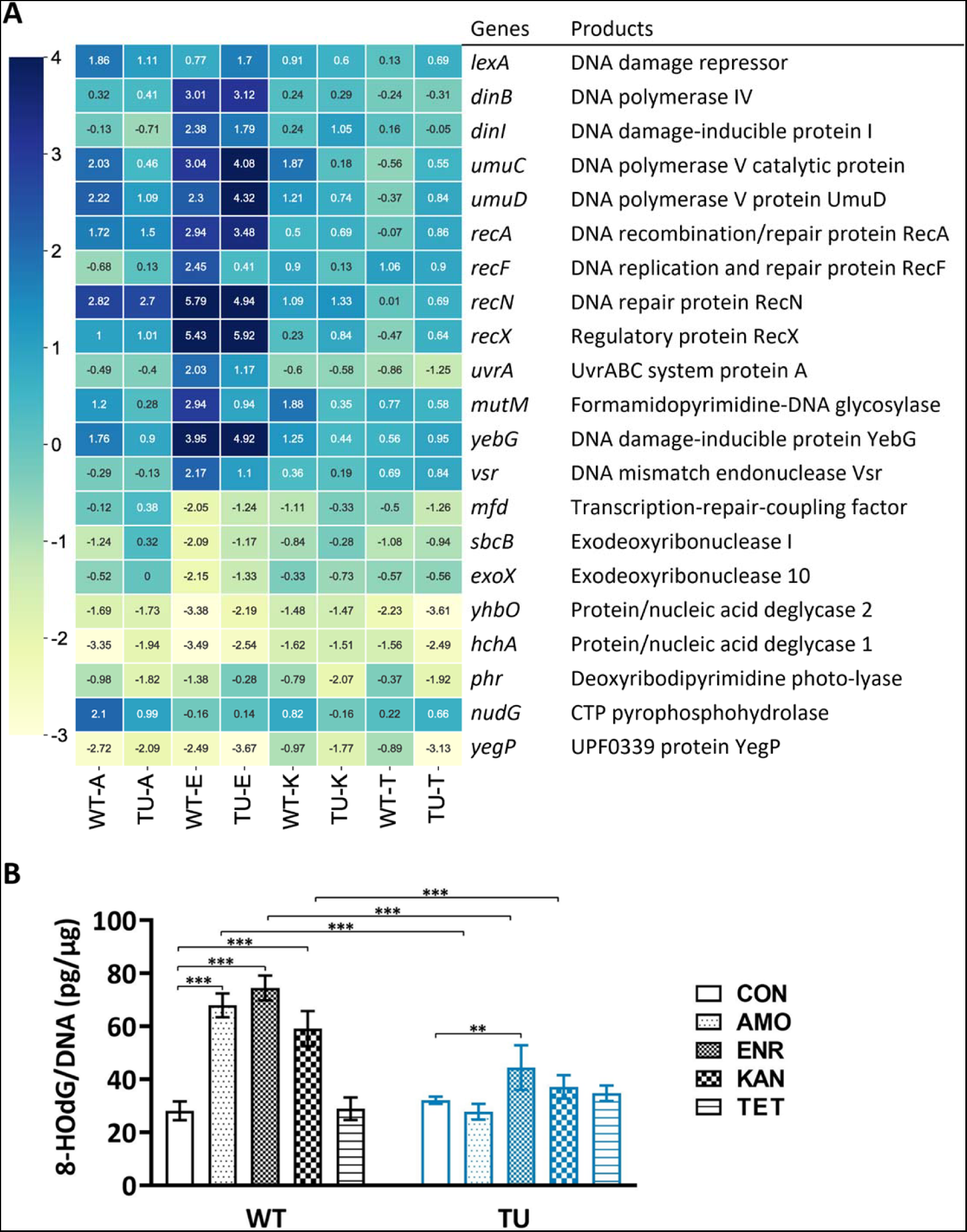
TU modulates SOS response genes, reducing *mutM* transcription as well as 8-HOdG levels after bactericidal antibiotics exposure. (A) Heat map showing DNA damage-repair-related gene expression Log_2_ fold change levels (blue, upregulated genes; yellow, downregulated genes). Rows show genes, columns indicate different strains. WT-A, resistance-evolved wild-type MG1655 treated with amoxicillin, TU-A, resistance-evolved TU-added MG1655 exposed to amoxicillin, etc., E, enrofloxacin, K, kanamycin, T, tetracycline. Scale bar showing the value of Log_2_ fold changes. (B) 8-HOdG production levels measured by DNA damage competitive ELISA. Culture used are referred to A. Data are represented as means ± SD, statistical significance was investigated using a one-way ANOVA, **p < 0.01, ***p < 0.001. CON, no-treated control, AMO, amoxicillin-treated, ENR, enrofloxacin-treated, KAN, kanamycin-treated, TET, tetracycline-treated.

**Figure 8.**
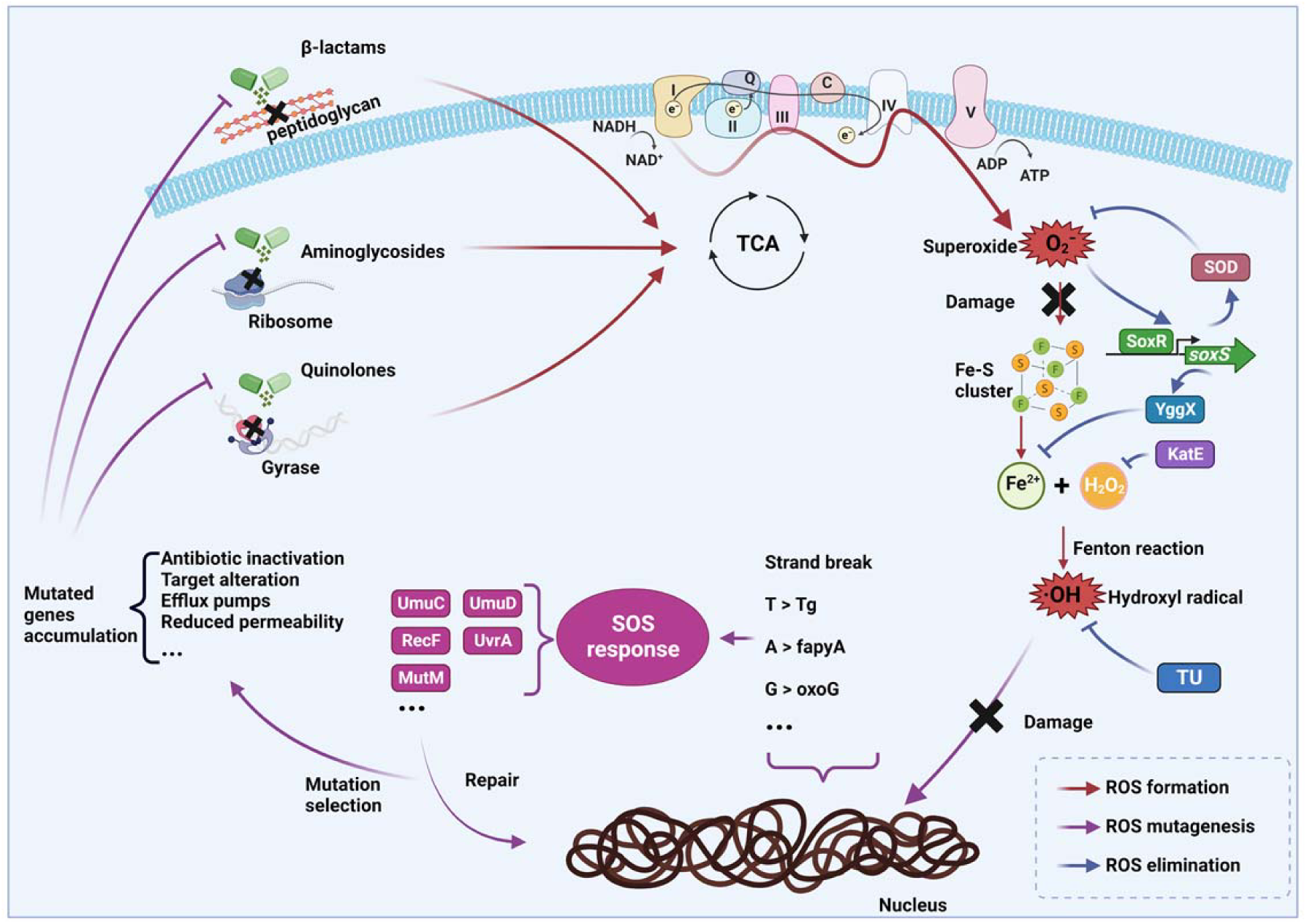
Scheme depicting the chain of events by which non-lethal concentrations of bactericidal antimicrobials cause mutations resulting in resistance to the specific antibiotic. In brief: TCA cycle activity mediated oxidation of NADH may lead to the emergence of superoxide causing damage to iron-sulfur cluster containing proteins which with hydrogen peroxide present can lead to the generation of the hydroxyl radical. The ensuing genome damage can promote the SOS response which, if initial levels of the reactive oxygen species (ROS) are sublethal, may promote mutation generation in surviving cells some of which are beneficial for antibiotic survival. The data presented are summarized by the processes and enzymes indicated and provide a mechanistic basis for the observed hormetic dose responses induced by antibiotics in bacteria (23).

To further clarify the emergence of the DNA damage biomolecule 8-HOdG, we measured its levels after antibiotic treatment. We choose the same strains as for RNA sequencing and performed the same treatment. In the TU absence group, the 8-HOdG levels were significantly increased after bactericidal antibiotics exposure (Figure 7B). Only enrofloxacin-treatment elevated the 8-HOdG levels in the TU-treated culture. Compared with the TU absence strains, the 8-HOdG levels in the TU treatment culture were significantly reduced. Together with the transcriptomics, we conclude that the ROS-induced DNA mutagenesis agent 8-HOdG was increased during the bactericidal antibiotics’ exposure and decreased after TU treatment.

## Discussion

The central question this study addressed is whether the ROS formed in *E. coli* upon exposure to bactericidal antibiotics has by the principle of hormesis a role in development of resistance. This hypothesis was postulated based on the capacity of ROS to induce DNA damage, such as the formation of 8-hydroxy-2’-deoxyguanosine (8-HOdG), which triggers the SOS response and activates the low-fidelity error-prone DNA repair systems, ultimately leading to increase the mutations accumulation. (41)(42). Two main observations support this notion 1) mutants that have reduced capabilities to eliminate ROS have higher rates of resistance development and 2) removal of ROS by thiourea decreased rates of resistance development. This suggests a mechanism for ROS-induced hormesis, killing occurs during short-time, high-dose antibiotic exposure, while the mutagenic effect during long-term, non-lethal dose exposure (43).

ROS can cause over one hundred different 2-deoxyribose modifications and oxidative nucleobase lesions (44). The electrophilic hydroxyl radical (•OH) can directly react with DNA nucleobases. For example, hydroxyl radicals attack the C5=C6 double bonds of thymine or cytosine (45). Fragmented formamidopyrimidine is formed by the hydroxyl radical-induced imidazole ring opening in guanine and adenine (46). Additionally, 8-HOdG is generated from hydroxylation of guanine in nucleotide pools or the genomic DNA level (47). Instead of pairing with cytosine, 8-HOdG prefers to pair with adenine thus causing frequent guanine-to-thymine mutations during replication(48), which were also observed in our experimental system (43). Consistent with this notion, the 8-HOdG level was increased after bactericidal antibiotics exposure, and it was significantly reduced after the application of ROS scavenger TU. These findings suggest that the emergence of ROS creates favorable conditions for elevated mutation rates. In addition to this direct nucleobase damage, hydroxyl radicals can interact with lipids, leading to the formation of malondialdehyde and 4-hydroxynonenal (49). These lipid peroxidation products can react with adenine, guanine, and cytosine to form mutagenic adducts (50). Along with the nucleobase damage, ROS can also compromise single-strand break and double-strand break (51)(52).

In response to ROS-induced DNA damage, bacteria activate SOS response genes, firstly *lexA* and *recA* (53). RecA searches for damaged DNA, while the breakdown of LexA activates SOS genes involved in damage repair (54). The activation of repair systems can exhibit a general nature or be specific to various antibiotics. During exposure to several antibiotics, the transcription levels of *recA* and *lexA* did not show an obvious common pattern between TU-treated or untreated conditions (Figure 7A). However, other damage-repair genes which are subsequently activated by the SOS response were upregulated by bactericidal antibiotic treatment, and this upregulation was attenuated by the addition of TU. For example, *umuC* and *umuD* upon amoxicillin or kanamycin exposure, *recF* and *uvrA* after enrofloxacin exposure. This indicates that transcription levels of DNA damage-repair genes differ as a result of exposure to different antibiotics and possibly of different ROS levels. Exposure to enrofloxacin yielded more upregulated genes. Conceivably because enrofloxacin targets DNA gyrase, and hence causes more direct DNA damage. Transcription levels of *mutM* were all upregulated upon exposure to the three bactericidal antibiotics and attenuated by TU treatment. *mutM* codes for a formamidopyrimidine-DNA glycosylase, with an inclination for, but not limited to, base excision repair for 8-HOdG. For example, it also recognizes and removes oxidized purines, and some oxidized pyrimidines to leave an apurinic or apyrimidinic (AP) site (55) (56). The transcription regulation of this glycosylase MutM characterizes the DNA damage-repair process that is activated upon exposure to bactericidal antibiotics. SOS response-induced genes, such as *umuC* and *umuD* which code for error-prone DNA polymerase V, contribute to an increase of mutation rates (57)(58). As a consequence, they might promote, at sublethal stress levels, the accumulation of beneficial antimicrobial resistance-conferring mutations (59)(60)(61) (62).

The low-fidelity polymerase V is derepressed when DNA is damaged. Due to the absence of an intrinsic 3’–5’ exonuclease proofreading activity, it is unable to rectify errors during translesion DNA synthesis (63). This error, however, creates the potential to overcome the DNA damage caused by ROS, thus producing damage-inducing mutations in the *E. coli* genome (64). During long-term sublethal levels of antibiotic exposure, mutations that favor resistance development will be selected from these damage-inducing mutations. This offers an opportunity for research on drugs that target the SOS response and so aim to reduce the development of drug resistance. For instance, N^6^-(1-naphthyl)-ADP prevents the formation of the RecA−DNA filament (65). Suramin inhibits the RecA-induced cleavage of LexA (66). Antioxidant drugs such as N-acetylcysteine can attenuate ROS levels, SOS response induction, and mutagenesis during ciprofloxacin treatment (67).

The various bacterial strategies for resistance development are reflected in the whole genome sequencing results. In general, these consist of pumping out, reducing entry, or degrading the drugs and of protecting, modifying, or changing the expression of drug targets (68). Although the strains at the end of each antibiotic treatment had evolved in a different manner, the final results showed commonality. The most characteristic mutations are in β-lactamase after amoxicillin treatment, DNA topoisomerase after enrofloxacin treatment, and elongation factor G after kanamycin treatment. *E. coli* increases the expression of β-lactamase by amplifying *ampC*, which is also evidenced by the fragments we observed containing *ampC* amplification (69). These contigs could come in different sizes, had copy numbers between 20 and 600, and were horizontally transferred, thereby making a sensitive *E. coli* receptor strain highly amoxicillin resistant (31)(70). Quinolone-induced mutations are often clustered within small regions of DNA topoisomerase-coding genes, known as quinolone resistance-determining regions (71). Mutations may lead to structural changes in the target site of the enzyme tetramer, resulting in reduced affinity for quinolones and thus resistance to these drugs (72). When *E. coli* is exposed to enrofloxacin, *gyrA* was first mutated, followed by *gyrB*, *parC*, or *parE* (35). Elongation factor G (EF-G) *fusA* catalyzes the GTP-dependent ribosomal translocation during translation elongation (73). Mutations in this gene may prevent kanamycin from binding EF-G and preventing translation. However, some mutations in *fusA* have showed several side-effects, including decreased levels of growth rate, reduced stringent response sensors (ppGpp), and increased sensitivity to oxidative stress (74)(75)(76). In addition to these target-specific mutations, other mutations, such as reduced membrane permeability and efflux pumps, appeared in combination. In this study, we also found mutations in many genes related to oxidative stress and SOS response. This is additional evidence that ROS and SOS response are involved in the evolution of drug resistance. Moreover, this explains why strains that had developed resistance to bactericidal antibiotics and were next exposed to other bactericidal antibiotics, showed increased rates of resistance acquisition (77). ROS and the subsequently activated SOS response accelerate resistance evolution in *E. coli* exposed to stepwise increasing sub-lethal levels of bactericidal antibiotics.

This study demonstrates the role of ROS in the development of antimicrobial resistance. Development of resistance in *E. coli* exposed to stepwise-increasing sub-lethal levels of bactericidal antibiotics, was slowed down by treatment with the antioxidant TU, and mutants with reduced ROS-scavenging showed accelerated resistance acquisition. However, no significant increase in the mutation rate was found in the mutant strains compared to the wild-type strains upon bactericidal antibiotics exposure which may well reflect that we have yet to pinpoint the exact concentration of ROS that is responsible for the tipping point from ROS induced cell death to ROS induced beneficial mutation acquisition. In future, we may strengthen the verification of our results by knocking out multiple genes or using the ROS-inducing chemicals such as Hydroxyurea to come closer to the indicated concentration domain (78). Considering that the purpose of our research is to understand the formation of resistance and try to reduce or slow down the resistance acquisition, our study demonstrates the feasibility of using antioxidants to reduce excess ROS formation in vivo and in that manner reduce drug resistance development by inhibiting SOS responses.

## Materials and Methods

### Bacterial strains, media, and growth conditions

The antibiotic sensitive *E. coli* K12 MG1655, and *E. coli* K12 Keio strains with specific gene deletions (Δ*sodA*, Δ*sodB*, Δ*soxR*, Δ*soxS*, Δ*katE*, and Δ*yggX*) were cultured in phosphate-buffered defined minimal Evans medium supplemented with 55mM glucose (pH 6.9) (79). When described “as TU-added” 100mM thiourea was added to the medium. Cultures were incubated in tubes with a start optical density (OD) 600nm of 0.1 at 37°C under constant shaking at 200 rpm. The kanamycin cassette in Keio strains was replaced by temperature-sensitive pCP20 plasmid.

### Evolution experiments

The evolution experiments aimed at inducing resistance were conducted in accordance with the previously established methodology (80). Briefly, susceptible naïve *E. coli* K12 strains of wild-type, TU-treated wild-type, and single-gene knockout strains (Δ*sodA*, Δ*sodB*, Δ*soxR*, Δ*soxS*, Δ*katE*, Δ*yggX*) were exposed to one-quarter MIC of amoxicillin, enrofloxacin, kanamycin, or tetracycline. An untreated control group without antibiotics was also included. After 24 hours of incubation, if the OD_600_ of the antibiotic-exposed culture was equal to or greater than 75% of the OD_600_ of the antibiotic-free culture, the antibiotic-exposed cells were selected and inoculated with double concentrations of the respective antibiotic, while retaining the previous low concentrations as a back-up. After another 24 hours of incubation, if the OD_600_ of the high antibiotic concentration culture was equal to or greater than 75% of the OD_600_ of the low antibiotic concentration culture, cells from the high concentration culture were used; otherwise, cells from the low concentration culture were used, and the antibiotic concentration was doubled again. The experiment was stopped when the bacteria could no longer tolerate the higher antibiotic concentration. Each strain’s resistance evolution was independently performed at least twice, and the antibiotic-free control groups were cultured in the absence of antibiotics but otherwise under identical conditions until the end of the experiment.

To monitor the development of resistance, MIC measurements were conducted three times a week. Cells were cultured in 96-well plates using a spectrophotometer plate reader (Thermo Fisher Scientific) with antibiotic concentrations ranging from 0.25 μg/mL to 2048 μg/mL, doubling at each step. The initial OD was 0.05 and the MIC was determined as the lowest concentration that resulted in a final OD_600_ of less than 0.2.

### ROS measurements

Fluorescence microscopy was performed to monitor ROS generation. The fluorescent dye 3′-(p-Hydroxyphenyl) Fluorescein (HPF) was utilized for the detection of ROS. Cells were sampled from cultures during the mid-stages of resistance development, specifically when they reached resistance concentrations of 32 μg/mL for each antibiotic. In each culture, HPF dye was added to achieve a final concentration of 10 μM, followed by incubation in a shaking incubator at 37°C for 40 minutes. Subsequently, bacterial cultures were centrifuged at 6000 rpm for 5 minutes, and the resulting pellet was thoroughly resuspended in medium. The samples were then treated with the corresponding antibiotic concentration, and 1.3 μL of each sample was pipetted onto a microscope slide glass with 2% agarose mixed with medium. Fluorescence detection was performed using the Nikon Eclipse Ti microscope equipped with NIS-elements AR software, employing an excitation/emission wavelength of 490/515 nm. The assembly of the time-lapse pictures was done using ImageJ software.

Flow cytometry (FACS) was used to quantify the level of ROS production. Overnight cultures were diluted to an OD_600_ of 0.1 in fresh medium, followed by treatment with the corresponding antibiotics. The cultures were then incubated for 4 hours at 37°C in a shaking incubator. Subsequently, a final concentration of 10 μM of the fluorescent dye HPF was added to each culture and incubated for an additional 40 minutes. All samples were centrifuged at 6000rpm for 5 minutes and the pellet was resuspended in 1 mL Evans medium without glucose. For the FACS sorting, BD FACSAria™ III Sorter was used with BD FACSDiva™ Software version 8.0.1. The laser settings were set on 250V FSC, 400V SSC, and 450V FITC (GFP). A total of 10,000 events were measured for each sample. The percentage of the ROS-positive population was determined, and each sample was independently replicated three times.

### Mutation rate measurements

Fluctuation assays were performed to determine the mutation rate, rifampicin was used to select for gain-of-function mutations in the *rpoB* gene (67). The experimental procedure was carried out as follows: Susceptible *E. coli* K12 strains, including wild-type, TU-treated wild-type, Δ*sodA*, and Δ*soxR* were exposed to one-quarter MIC of amoxicillin, enrofloxacin, kanamycin, and tetracycline. Each culture was incubated until reaching OD_600_ of approximately 0.5 to 0.6. Next, 200 µL of each culture was plated onto LB agar plates containing rifampicin (50 µg/mL), while 100 µL of each culture was stepwise diluted to 10^-7^ using medium and plated onto antibiotic-free LB agar plates. The plates were then incubated at 37°C for 24 hours for the antibiotic-free LB plates and 48 hours for the LB plates containing rifampicin. Colony-forming units (CFUs) were counted for both plate types. Mutation rates were calculated by Lea-Coulson Ɛ < 1.0, the comparison was done by the maximum likelihood ratio statistical test in webSalvador 0.1, which was powered by rSalvador 1.8 (30). WebSalvador provided the maximum likelihood ratio value, along with the corresponding P-value and the upper and lower limits of the 95% confidence intervals, which were used to generate the bar graph. Statistical significance was indicated as *p < 0.05 and **p < 0.01. The experiment was independently replicated at least 10 times to ensure the reliability and reproducibility of the results.

### Whole genome sequencing

The genomic DNA was extracted from the final resistant strains and the corresponding antibiotic-free strains utilizing the DNeasy Blood and Tissue Kit (Qiagen). Libraries were constructed using the NEBNext Ultra II FS DNA Library Prep Kit for Illumina (New England BioLabs) in combination with NEBNext Multiplex Oligos for Illumina (96 Unique Dual Index Primer Pairs; New England BioLabs), following the manufacturers’ recommended protocols. Subsequently, the genomic DNA libraries were subjected to paired-end sequencing (2 x 150 bp) on the NextSeq 550 next-generation sequencing system (Illumina). The quality of the raw reads was assessed using FastQC and MultiQC. Adapter sequences were removed using Cutadapt. After removing the low-quality bases and optical duplicates, the reads were aligned to reference genomes of wild-type strains (NC000913) and Keio strains (CP009273) using Bowtie2, and the PCR duplicates were removed. Variant calling was performed using Freebayes and Lofreq, and variant annotation was conducted using Snpeff. Subsequently, the single nucleotide polymorphisms (SNPs) and small indels were inspected using the Integrative Genomics Viewer (IGV). The mutated genes in the resistant strains were compared to their corresponding control (antibiotic-free) strains, and shared mutated genes were excluded from further interpretation. In addition to analyzing small genomic alterations, copy number analysis by cn.MOPS was performed to identify larger genomic alterations, such as amplifications and deletions.

### RNA sequencing

Total RNA was isolated and purified from the final stable resistant strains and their corresponding antibiotic-free strains using the RNeasy Protect Bacteria Kit (Qiagen). For RNA-Seq analysis, libraries were constructed following the manufacturer’s protocols, employing the NEBNext rRNA Depletion Kit (Bacteria) (New England BioLabs) in combination with the NEBNext Ultra II Directional RNA Library Prep Kit for Illumina and NEBNext Multiplex Oligos for Illumina (Unique Dual Index Primer Pairs) (New England BioLabs). The resulting libraries were subjected to sequencing on a NextSeq 550 Sequencing System (Illumina) with read lengths of 75 bp. To ensure data quality, FastQC and MultiQC were employed for comprehensive quality control of the raw sequencing data. Subsequently, the reads underwent trimming procedures, and were aligned to the reference genomes (NC000913) using Bowtie2. For the determination of differential gene expression, normalized gene expression values were computed, and Log_2_ fold changes were calculated by comparing the resistant strains to the antibiotic-free control by HTSeq and DESeq2. Within each treatment group, DNA damage-repair genes surpassing a cutoff of 2 were selected and incorporated into the generated heat map.

## 8-HOdG level measurements

The DNA Damage Competitive ELISA Kit was employed to quantify the levels of the oxidative stress marker 8-hydroxy-2’-deoxyguanosine (8-HOdG) according to the manufacturer’s instructions. In brief, the final stable resistant strains of wild-type and TU-treated wild-type were selected and subjected to treatment with the corresponding antibiotics. The cultures were incubated for 4 hours at 37°C in a shaking incubator. Subsequently, the samples were centrifuged at 6000rpm for 5 minutes, and the resulting pellet was resuspended in 0.3 mL of Lysis Buffer (containing 10 mM Tris-HCl, 2 mM EDTA, 1% SDS). A 5-fold dilution of the samples was loaded onto the antibody-coated 96-well plate. Following the recommended protocol for incubation, the absorbance at 450 nm was measured using a spectrophotometer plate reader (Thermo Fisher Scientific). The concentrations of 8-HOdG in the samples were determined by referencing to a standard curve. Furthermore, the DNA concentrations of the samples were measured using a microvolume spectrophotometer (DeNovix) and utilized for normalization. As controls, antibiotic-free samples of wild-type and TU-treated wild-type were included, and each sample was independently replicated three times.

### Quantification and statistical analysis

Statistical analyses were conducted using IBM SPSS Statistics software. Detailed information regarding the statistical methods employed for each experiment can be found in the figure legends and corresponding figures.

### Data availability

The binary alignment/map (bam) files of the sequenced strains have been deposited in the NCBI database and can be accessed at BioProject PRJNA987436 (FACS), PRJNA954686 (WT), PRJNA987546 (TU), PRJNA987571 (Δ*sodA*), PRJNA987595 (Δ*sodB*), PRJNA987616 (Δ*soxR*), PRJNA987630 (Δ*soxS*), PRJNA987644 (Δ*katE*), PRJNA987659 (Δ*yggX*), and PRJNA988039 (RNAseq).

## Acknowledgments

We thank the Van Leeuwenhoek Center for Advanced Microscopy (LCAM) at the University of Amsterdam for offering the microscopy platform. We thank Selina van Leeuwen for her help with the sequencing works. The students Maria Mazmanidou, Mark Kok, Reina Groot, and Mireia Novell Cardona performed experiments as part of their degree requirements.

This study was financed by the Netherlands Food and Consumer Product Safety Authority (NVWA). The NVWA was not involved in design of the experiments, analysis of the data or writing the manuscript.

## Author contributions

WQ and BtK conceived the project. SB assisted in the design of experiments. WQ performed experiments and analysis of the data. MJ and WL performed the bioinformatic analysis. WQ and BtK wrote the manuscript. All authors critically reviewed the manuscript and approved the final version.

## Supplemental information

Table S1 Gene mutations in ROS positive and negative populations

Table S2 Gene mutations in antibiotic resistant strains

